# Transcriptome Analysis Reveals Long-Term Somatic Memory of Stress in The Woody Perennial Crop, Grapevine *Vitis Vinifera* cv. Cabernet Sauvignon

**DOI:** 10.1101/2024.04.06.588326

**Authors:** Jia W. Tan, Kiflu Tesfamicael, Yikang Hu, Harshraj Shinde, Everard J. Edwards, Penny Tricker, Carlos Marcelino Rodriguez Lopez

## Abstract

Plants can generate a molecular memory of stress resulting in primed plants that are more resilient to subsequent stresses occurring days to weeks after the priming event. Whether such a priming effect is maintained over longer periods, and after winter dormancy in perennial plants, is less studied. Here, we used whole transcriptome and methylome sequencing of grapevine plants over two growing seasons to characterize grapevines’ response to combined drought and heat stress in naïve and primed plants. Our results showed changes in expression of genes associated with epigenetic modifications during stress and after stress removal, suggesting the establishment of epigenetic memory of stress. Primed plants had a small number of differentially expressed genes associated with stress response one year after the priming event even in the absence of second stress and presented a stronger transcriptional response than naïve plants when re-exposed to stress. Methylome analysis revealed an increase in DNA methylation in primed vines under combined stress, and that methylation patterns were less variable among plants under stress than control plants. We did not observe any correlation between DNA methylation and gene transcription, suggesting that stress-induced expression changes were, at least partially, independent of DNA methylation, with posttranscriptional regulation and histone modifications more likely candidates in the establishment of epigenetic memory. Additionally, we characterized stress responsive genes based on their transcriptional profile and function and propose a new comprehensive and intuitive classification model for stress memory genes in perennials.

## Introduction

Viticulture is highly dependent upon climatic conditions during the growing season. Climate determines the suitability to grow a particular variety, as the most desirable composition of grapes requires specific climatic conditions (Gladstones, 1992). Heat and drought are common abiotic stress factors often connected to grapevine yield losses (Vinocur and Altman, 2005). Although normally studied in isolation, such losses often result from both stresses acting in combination (Vogel et al., 2019). The progression of grapevine transcriptional responses to acute combined heat and drought stress in one season has been reported (Tan et al., 2023), but chronic and recurring stresses in different seasons are often observed in nature (Pagay et al., 2022), and responses to recurring stress are much less understood.

Growth conditions that inhibit normal growth and development can trigger a priming response in plants. Priming can be defined as a modified response when a plant is re-exposed to stress compared with that of a naïve plant (unprimed) (Aranega-Bou et al., 2014). In general, priming is evidenced by positive effects like a stronger or faster response pattern (Bruce et al., 2007; Conrath, 2009; Crisp et al., 2016). The maintenance of this memory can be somatic (i.e., transmitted by somatic cells within the plant exposed to the stress) or inter- or transgenerational stress memory (transmitted to the offspring via the germline of the plant exposed to the stress) (Lämke and Bäurle, 2017). Molecular mechanisms for the storage and retrieval of this stress memory may include epigenetic regulation, transcriptional priming, the primed conformation of proteins, or specific hormonal or metabolic signatures (Crisp et al., 2016; Ding et al., 2012; Hake and Romeis, 2019; He and Li, 2018; Heil and Karban, 2010). Evidence suggests that stress memory is heavily epigenetically-based and involves mechanisms such as chromatin remodeling, DNA methylation, nucleosome position, histone modification, and noncoding RNA-mediated regulation (Liu et al., 2021). It is believed that stress-induced epigenetic marks are the molecular base for long-term and transgenerational maintenance of priming (Tricker et al., 2013a), and that this stress memory can be observed through the physiological, transcriptional, and biochemical modifications occurring when exposed to a stress factor in the future, so that the plant has become more resistant, or sensitive, to the same (Alves de Freitas Guedes et al., 2019; Perrone and Martinelli, 2020) or different stress (Tricker et al., 2013b). The duration of stress memory will depend on the stability of the epialleles responsible for the stress memory, either mitotically or meiotically. In mitotically stable memory, it has been observed that plant epigenetic (e.g., DNA methylation) profiles are predictive of the environment where the plant grows (Xie et al., 2017), and that such changes are persistent during vegetative growth, throughout newly developing tissues, and through the lifetime of the plant (Deleris et al., 2016; Lämke and Bäurle, 2017).

In transcriptomes, priming can be evident as a change in the expression of certain genes in primed plants when exposed to a second stress. Stress responsive genes can be classified as: non-memory genes (i.e., those in which expression is the same in primed and naïve plants when exposed to stress); and memory genes (i.e., those in which expression is significantly different in primed and naïve plants). Two main memory gene classification systems have been proposed to date. Ding et al. (2014) defined six types of memory genes, i.e., (+/+), (-/-), (+/-), (-/+), (+/=), and (-/=); where the first symbol indicates the direction of the transcriptional changes occurring in plants exposed for the first time to stress compared to control plants (+ and – indicate an increase or decrease in expression of a given gene respectively), and the second symbol indicates the transcriptional changes of a primed plant compared to its naïve state response. Bäurle (2018), proposed a simpler classification system with non-memory genes (as defined above), and type I and type II memory genes. Type I genes maintain the direction (upregulation or downregulation) of alteration in transcription past the duration of the priming environmental stressor, while Type II genes present a modified response in expression after the triggering stress compared to the priming stress, following a lag phase of transcriptional inactivity. Although both models are complementary, they both fail to capture all possible types of memory genes (e.g., Ding et al. do not include Type I genes, while Bäurle does not describe Type I gene expression patterns in response to a triggering stress).

The majority of the studies of this priming effect or the memory of stress have been conducted in annual/model plants such as Arabidopsis (e.g., Ding et al., 2012). How the memory of stress is maintained in perennial plants after winter dormancy is less studied. A few studies of stress memories in perennials exist for: coffee plants (*Coffea canephora*) (de Freitas Guedes et al., 2018), wild strawberries (*Fragaria vesca*) (López et al., 2022) and the perennial grass species tall fescue (*Festuca arundinacea*) (Bi et al., 2021). However, none of these studies covered more than one year of winter dormancy. Grapevine has recently been proposed as a model plant to study epigenomics in perennial plants due to its characteristics (Fortes and Gallusci, 2017) such that grape flower development is programmed one year in advance and that the environmental conditions of the previous year affect flower and subsequent fruit development, suggesting that a memory of the environmental conditions is established every year in meristems committed to flowering. Therefore, this makes grapevine an interesting model to study how long-term somatic stress memory is maintained after winter dormancy (Tan et al., 2023a).

Multi-omics approaches such as transcriptomics, epigenomics, degradomics, proteomics, and metabolomics have been developed and deployed to study the mechanistic basis of plant stress memory (Liu et al., 2021). In this study, we used transcriptome and methylome sequencing to study the potential role of epigenetic regulation during stress response, stress memory establishment, and the maintenance of long-term (two growing seasons) somatic memory in grapevine; and to identify and characterize the expression patterns of genes associated to somatic memory of stress in grapevine.

## Results

### Gene expression analysis

Transcriptome sequencing data de-multiplexing yielded an average of 25 million reads per sample after quality filtering (QC 30). The average percentage of mappable reads per sample was 82%, ranging from 70-92% (Table S1).

#### Identification of modified responses in gene expression as a result of priming

First the gene expression of ST4_2_00 and ST4_2_03 plants was compared to identify the genes differentially expressed under combined stress. This comparison served two functions, first to identify the genes differentially expressed by naïve plants when exposed to a first stress event. Secondly, these results also served as validation of the results presented by Tan et al. (2023b) for season 1, as the naïve plants priming state and growing conditions replicated those in that experiment. In this comparison, 176 genes were found to be up-regulated, and 431 were down-regulated (Figure 1A) in naïve plants grown under stress conditions (ST4_2_03), compared to naïve plants grown under control conditions (ST4_2_00). Pathway analysis revealed pathway enrichment similar to those during the season 1 experiment, such as ‘plant hormone signal transduction’, ‘protein processing in endoplasmic reticulum’, and ‘phenylpropanoid biosynthesis’ (Figure S1).

**Figure 1.**
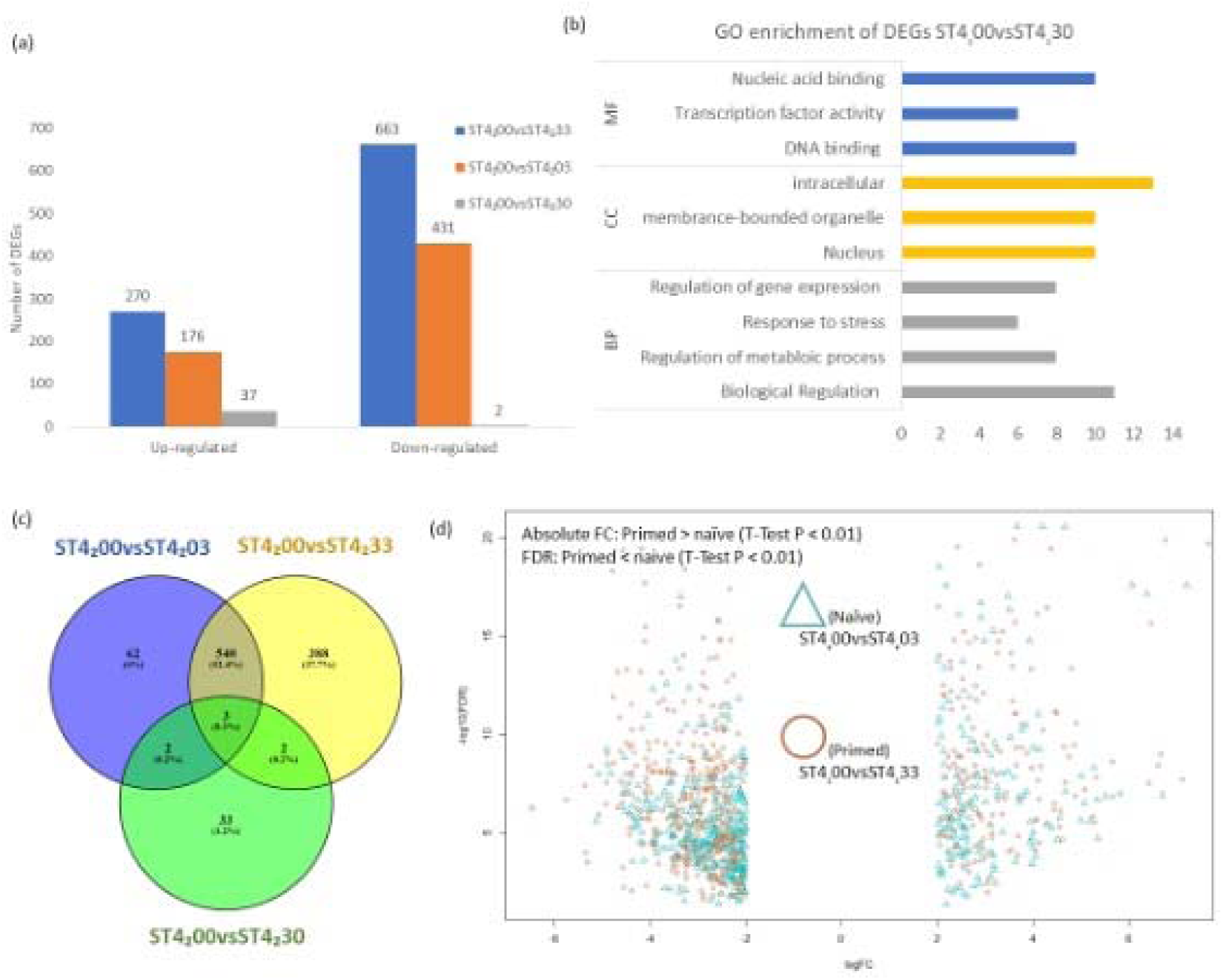
Analysis of differential gene expression between naïve and primed plants under stress or control conditions. (a) Number of differentially expressed genes (DEGs) identified between naïve plants grown under control conditions on year 2 (ST4_2_00) and: primed plants exposed to stress on year 2 (ST4_2_33), naïve plants exposed to stress on year 2 (ST4_2_03) and primed plants exposed to control conditions on year 2 (ST4_2_30). (b) Significantly enriched GO terms for DEGs identified in ST4_2_00vsST4_2_30. (c) Common and unique DEGs for each of the comparisons described above. (d) Effect of priming on the magnitude of change in expression (i.e., log fold change (horizontal axis) and FDR (vertical axis) of the common 543 DEGs identified in primed and naïve plants exposed to stress on year 2. Brown circles represent the common DEGs found in ST4_2_00 (naïve plants under control conditions) compared to ST4_2_33 (primed plants under combined stress). Blue triangles represent the common DEGs found in ST4_2_00 compared to ST4_2_03 (naïve plants under combined stress).

The existence of an epigenetic memory in plants exposed to stress in season 1 was then assessed using two different comparisons: First, the gene expression of primed plants was analyzed in the absence of a triggering stress event by determining differential gene expression between ST4_2_00 vs ST4_2_30 plants (i.e., naïve and primed plants in the absence of recurring stress respectively). In this comparison, 37 genes were found to be up-regulated, and 2 down-regulated in primed plants compared to naïve ones (Figure 1A). A histone lysine N-methyltransferase ATXR6-associated gene was found to be up-regulated among the identified DEGs. Other DEGs identified included APETALA 2/Ethylene Responsive Factor (AP2/ERF), no apical meristem, *Arabidopsis thaliana* activating factor and cup-shaped cotyledon (NAC), WRKY, ATP-binding cassette (ABC) transcription factor family genes, as well as F-box domain-containing and cysteine-rich transmembrane module (CYSTM) domain-containing proteins (Table S2). GO analysis of identified DEGs revealed 57 significantly enriched GO terms (Supplementary Table S2), with the top molecular function and biological process GO terms including nucleic acid/DNA binding, transcription factor activity, regulation of gene expression, and ‘response to stress’ (Figure 1B). Second, the number of DEGs in primed plants under a repeated stress event (i.e., ST4_2_00 vs ST4_2_33) was calculated and compared to those identified above in naïve plants exposed to stress in season 2 (i.e., ST4_2_00 vs ST4_2_03). Although the majority of DEGs found in both types of plants (543) were found to be commonly regulated, that is, they were up-or down-regulated in both primed and naïve plants (Figure 1C), primed plants showed a higher number of unique DEGs than naïve plants exposed to stress for the first time (i.e., 390 vs 64 DEGs respectively (Figure 1C)). GO analysis of naïve plant exclusive DEGs showed enrichment in histone methylation, covalent chromatin modification, and histone lysine methylation (Table S3A). Similar GO terms were observed in first-year plants by Tan et al. (2023b) under combined treatment. DEGs exclusive to primed plants exposed to a second stress were enriched for anatomical structure development and developmental process and GO terms associated with methylation were also identified (Table S3B). The GO analysis of the 543 DEGs identified in naïve and primed plants exposed to stress showed enrichment of chromatin assembly, methylation, and chromatin organization (Table S3C).

To better understand the difference between naïve and primed plant responses to stress, and the effect of priming on gene expression, the magnitude of the expression changes in DEGs common between primed and naïve plants was compared (Figure 1D). For this, paired T-tests were performed for the individual fold changes (FC) of upregulated and down regulated genes, and the false discovery rates (FDR) of those genes (n=534). This analysis indicated that the fold change in expression of DEGs common to primed and naïve plants was larger and more significant (FDR) in primed plants (T-Test -FC and +FC, p < 0.01; T-Test FDR, p < 0.01).

#### Identification of putative stress memory genes

Memory genes are traditionally defined as those in which response is different in primed than in naïve plants. To identify putative memory genes in grapevine, we used gene expression clustering analysis on combined stress-induced DEGs identified in year 1 (that is DEGs between control and stressed plants identified at season 1 sampling times ST4_1_3 (671 DEGs) and ST5_1_3 (224 DEGs) (ST4 and ST5 hereafter for simplicity). ST4 DEGs formed 10 clusters (C0_ST4_ to C9_ST4_) containing a total of 384 genes (Figure S2), and ST5 DEGs formed 5 clusters (C0_ST5_ to C4 _ST5_) containing 101 genes (Figure S3). Among those, two clusters (C0_ST4_: 96 genes, and C6_ST4_: 20 genes) contained genes with similar expression levels in response to the priming and the triggering stress with an intermediate phase of no transcriptional differences (compared to the control plants) between stresses and were deemed non-memory genes (Figure 2 and Figure S2). Nine clusters (C1_ST4_: 21 genes; C3_ST4_: 57 genes; C4_ST4_: 26 genes; C7_ST4_: 20 genes; C8_ST4_: 71 genes; C0_ST5_: 20 genes; C2_ST5_: 12 genes; C3_ST5_: 14 genes; and C4_ST5_: 34 genes) (Figure S2, & S3) contained genes in which expression was maintained at significantly different levels from the control plants (ST4_1_0) during the time between the removal of the priming stress, and the triggering stress (i.e., ST6), so were deemed Type I memory genes. These clusters could further be divided into four different subgroups in relation to their response to the triggering stress. Genes in clusters C4_ST5_ and C7_ST4_ presented the same level of expression change in response to the priming and the triggering stress (Type I^=^) (Figure 2). Genes in cluster C0_ST5_ showed a significantly higher change in expression to the priming than the triggering stress (Type I^+^) (Figure 2). Genes in clusters C1_ST4,_ C3_ST4,_ C4_ST4,_ and C3_ST5_ showed a significantly lower change in expression to the priming than the triggering stress (Type I^-^) (Figure 2). All these clusters presented a change in expression between physiological recovery to the priming stress and the triggering stress. Conversely, no change in expression was observed in response to the triggering stress in clusters C2_ST5_ and C8_ST4_ when compared to plants at physiological recovery (Type I^0^) (Figure 2).

**Figure 2.**
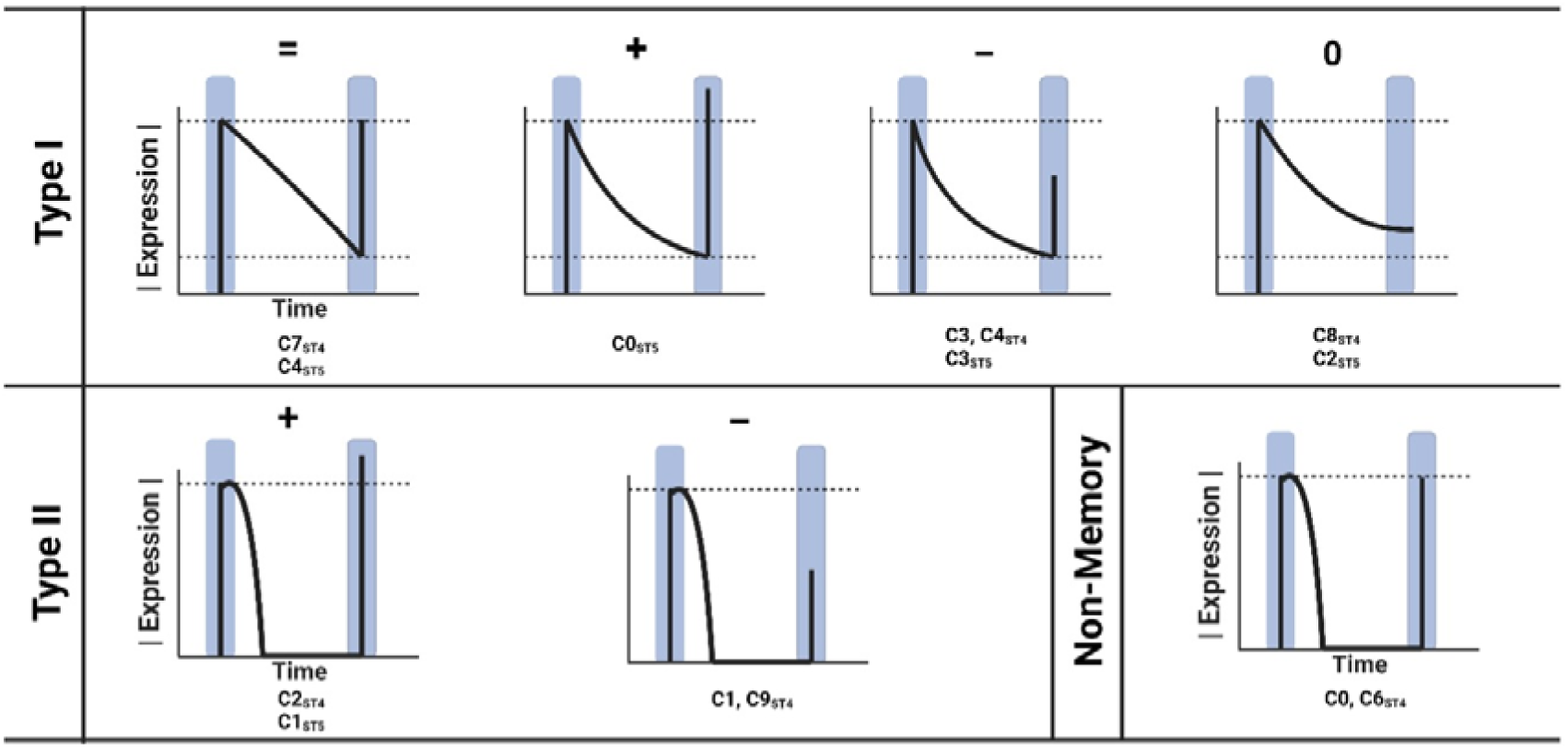
Stress memory gene models based on the expression patterns of DEGs found over two growing seasons. Type I: genes that exhibited sustain expression after first stress encounter. Type I^=^: expression changes upon second stress encounter was highly identical to the first. Type I^+^: expression changes upon second stress encounter was higher than the first. Type I^-^: expression changes upon second stress encounter was lower than the first. Type I^0^: the expression changes upon second stress encounter was minimal compared to the sustained expression after the first stress encounter. Type II: genes that exhibited expression changes between first and second stress encounter, the expression level returned to basal after the first stress. Type II^+^: the expression level of genes upon second stress encounter was higher than the first. Type II^-^: the expression level of genes upon second stress encounter was lower than the first. Non-memory: genes that exhibited no expression changes between first and second stress encounter, the expression level returned to basal after the first stress.

Three clusters (C2_ST4_: 28 genes; C9_ST4_: 26 genes; and C1_ST5_: 21 genes) contained genes presenting a modified response to the triggering stress compared to the priming stress, following a lag phase of transcriptional inactivity, and so, were deemed Type II genes. As with Type I genes, these clusters separated into different subtypes. C2_ST4_ and C1_ST5_ presented an enhanced change in expression to the triggering stress compared to the priming stress (Type II^+^) (Figure 2). Finally, C9_ST4_ genes presented a diminished change in expression to the triggering stress compared to the priming stress (Type II^-^) (Figure 2).

GO and functional analysis of all clusters containing memory genes as defined above was performed (Table S4). Interestingly, the GO terms for 21 genes from cluster C1 of ST5 DEGs were enriched in methylation, including histone H3-K9 methylation and DNA methylation. Among those, a structural maintenance of chromosome protein-associated gene was identified through functional annotation.

### DNA methylation analysis

An average of 69 million reads per sample were produced from the EM-seq library after quality filtering. The average percentage of mappable reads per sample to the PN40024 v.4 genomes was 54%. The average non-bisulfite conversion rate among the samples was 0.2%, and the average bisulfite conversion rate among the samples was 95.2%. The average percentage of covered bases was 81.22%, while the sequencing depth was 17X per sample (Table S5).

#### Global DNA methylation pattern induced by repeated combined stress in Grapevine

Analysis of the average methylation percentage (methylated cytosines, mCs) for each of the three contexts (CG, CHG, and CHH) showed that the CG context was the more methylated of the three, followed by CHG, and finally CHH (Figure 3A). Both naïve plants under stress (ST4_2_03), and primed plants in control conditions (ST4_2_30) showed similar context-specific DNA methylation to naïve plants under control conditions (ST4_2_00). Conversely, primed plants under stress conditions (ST4_2_33), presented a significant increase in mCG, mCHG, and mCHH (T-test, p ≤ 0.05) (Figure 3A). The PCA plot suggested that the global DNA methylation pattern in naïve plants (ST4_2_00 & ST4_2_03) was more variable compared to the primed plants (ST4_2_30 & ST4_2_33) (Figure S4).

**Figure 3.**
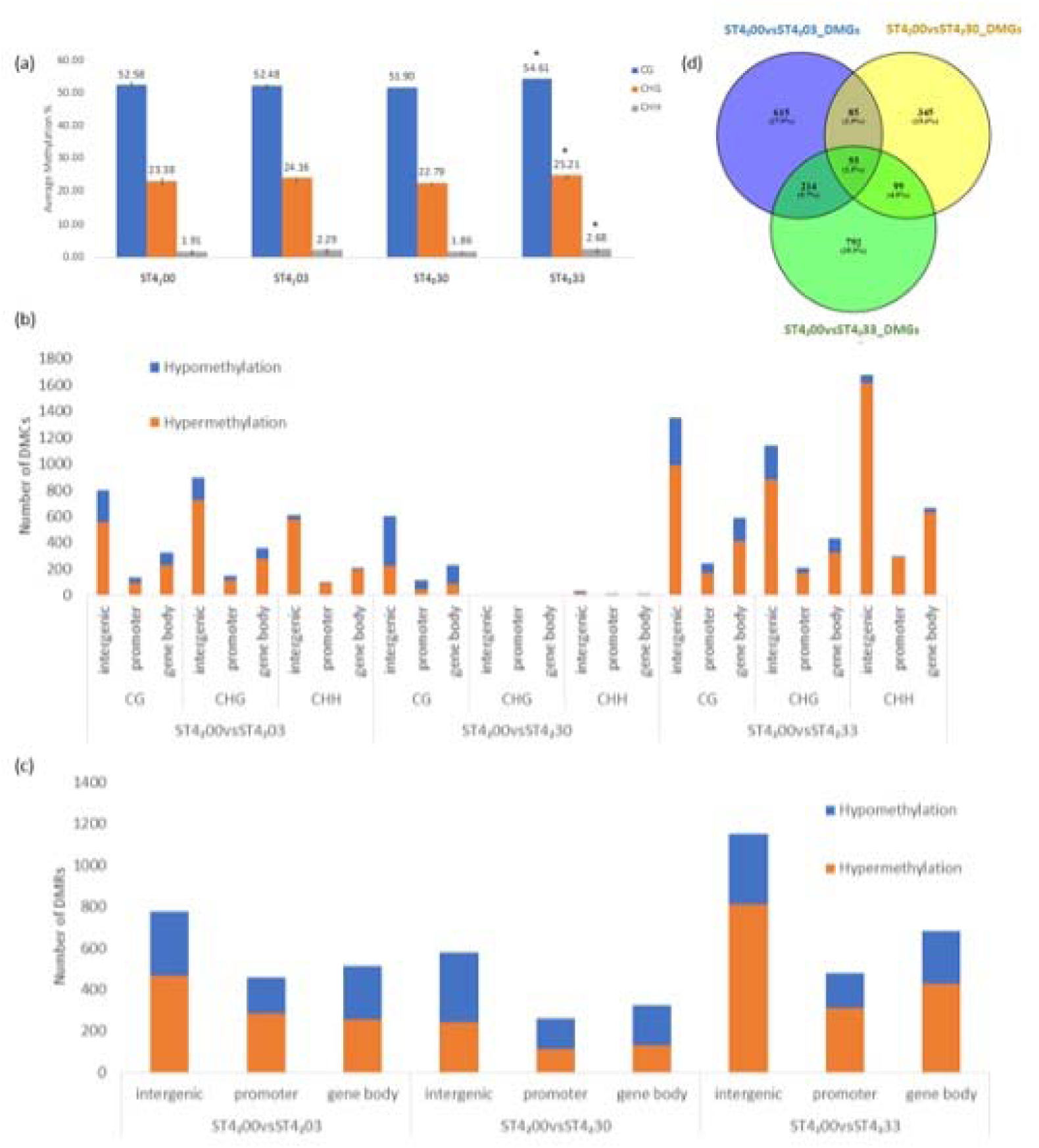
Effect of combined drought and heat priming and triggering stresses on grapevine DNA methylation. (a) Average DNA methylation level for each cytosine context (CG, CHG, CHH) between plant groups: ST4_2_33 (primed plants under combined stress), ST4_2_03 (naïve plants under combined stress) and ST4_2_30 (primed plants under control conditions) and ST4_2_00 (naïve plants under control conditions) asterisks indicates the significance (Student’s T test, p ≤ 0.05) of the difference between ST4_2_33, ST4_2_03, and ST4_2_30 compared to ST4_2_00. (b) Number of hyper-(hyper-DMCs) and hypomethylated differentially methylated cytosines (hypoDMCs) separated by sequence context and group comparison. (c) distribution of hyper-(hyperDMRs) and hypo-methylated differentially methylated regions (hypoDMRs) and in genomic features: promoter, gene body, and intergenic regions. (d) Venn diagram of differentially methylated genes between different plant groups.

The effect that priming and of stress in naïve and primed plants on local DNA methylation was determined by identifying differentially methylated cytosines (DMCs) and differentially methylated regions (DMRs) in the following comparisons ST4_2_00 vs ST4_2_30, ST4_2_ 00vs ST4_2_03, ST4_2_00 vs ST4_2_33. Briefly, plants exposed to stress for the first time presented a higher number of DMCs (1,254, 1,395, and 905 DMCs for CG, CHG, and CHH contexts, respectively), than primed plants in the absence of a triggering stress (938, 8, and 52 DMCs in CG, CHG, and CHH contexts, respectively), while primed plants under the effect of a triggering stress had the largest number of DMCs in all three contexts (2,178, 1,779, and 2,637 for CG, CHG, and CHH, respectively) (Figure 3B). The majority of DMCs were found in intergenic regions, regardless of context and comparison, and DMCs were more likely to be hypermethylated than hypomethylated (Figure 3B). The number of differentially methylated regions (DMRs) was assessed to study the dynamics of DNA methylation at specific loci. As with DMCs, the total number of DMRs observed ranked from ST4_2_00 vs ST4_2_33 (2,312 DMRs), ST4_2_00 vs ST4_2_03 (1,749 DMRs), to ST4_2_00 vs ST4_2_30 (1,161 DMRs) (Figure 3C). Also, as in the pattern observed for DMCs, the majority of DMRs identified were intergenic regions (55-57%), followed by gene body (25-27%) and promoter (17-19%), and were more likely to be hypermethylated than hypomethylated, except the DMRs in intergenic and promoter regions for ST4_2_00 vs ST4_2_30 where more DMRs were hypomethylated (Figure 3C).

As seen with DMCs and DMRs, the number of genes overlapping with a DMR (DMGs hereafter) was higher in primed plants under a triggering stress (ST4_2_33-DMGs = 1160), followed by naïve plants under stress (ST4_2_03-DMG = 969), then primed plants in the absence of a triggering stress (ST4_2_30-DMG = 584). Comparison of all DMGs identified showed that most were unique to each of the conditions (ST4_2_30, 03, and 33), while only 2.5% were common to all three conditions, 12.2% where common to ST4_2_03 and ST4_2_33 plants, and 7% to ST4_2_30 and ST4_2_33 plants (Figure 3D). The magnitude of the methylation changes in DMGs common between primed and naïve plants was then compared (269 DMGs). Unlike for gene expression in primed plants, there was no significant difference in level of changes for methylation between common DEGs in primed and naïve plants under stress.

GO analysis performed on DMGs in naïve plants exposed to stress (ST4_2_03) showed similar enrichment terms regardless of methylation change pattern (hyper- or hypo-methylated), such as ‘developmental process’, ‘protein serine/threonine kinase activity’, ‘reproduction’, and ‘response to stress’ (Table S6A-B). Whereas primed plants in the absence of a triggering stress (ST4_2_30) revealed a significant enrichment in GO terms such as ‘transcription factor activity’ and ‘histone modification’ both for hyper and hypomethylated DMGs. The term ‘signal transduction’ was unique to hyperDMGs, while ‘pyrophosphatase activity’ and ‘post-transcriptional regulation of gene expression’ were unique to hypoDMGs (Table S6C-D). Finally, primed plants under a triggering stress event (ST4_2_33) showed significantly enriched GO terms such as ‘response to stress’, ‘chromatin modification’, and ‘gene silencing’. Terms ‘mRNA metabolic process’ and ‘protein modification by small protein removal’ were unique to hyperDMGs and hypoDMGs, respectively. (Table S6E-F)

Changes in gene methylation over time were examined by comparing DMGs identified in plants under combined stress (ST4_1_3), and during physiological recovery (ST6_1_3) in season 1, and in primed plants under a triggering stress (ST4_2_33) (Figure 4A). 20 genes were differentially methylated at all three time points. Among these, 4 were hypomethylated at all three time points, 2 were hypomethylated in ST4_1_3 and ST6_1_3 but hypermethylated in ST4_2_33. 2 were hypomethylated in ST4_1_3 and ST4_2_33 but hypermethylated in ST6_1_3, 7 were hypomethylated in ST4_1_3 but hypermethylated in both ST6_1_3 and ST4_2_33, 2 were hypermethylated in both ST4_1_3 and ST4_2_33 but hypomethylated in ST6_1_3, 1 was hypermethylated in ST4_1_3 but hypomethylated in ST6_1_3 and ST4_2_33. Lastly, 2 were hypermethylated in ST4_1_3 and ST6_1_3 but hypomethylated in ST4_2_33 (Figure 4B).

**Figure 4.**
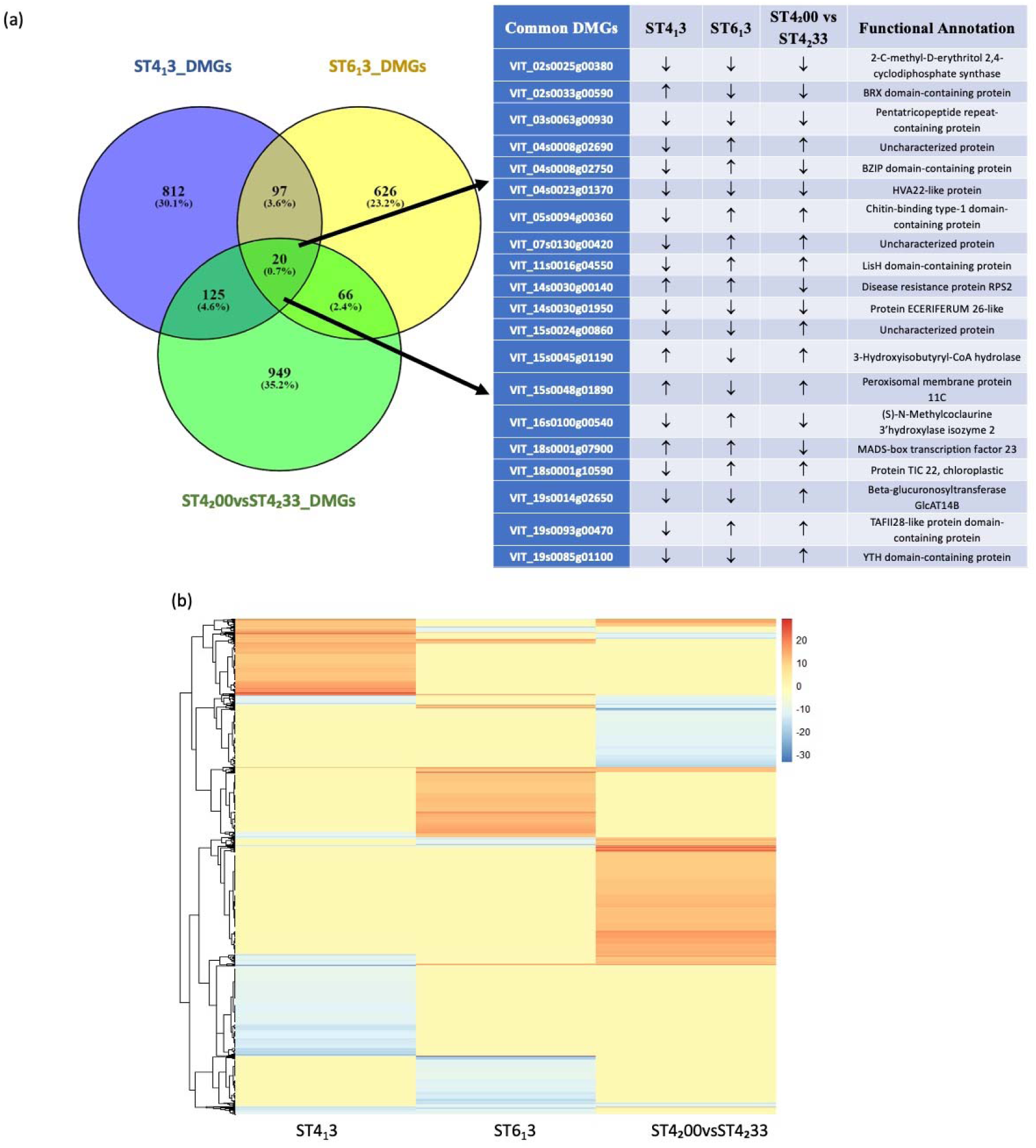
Changes in gene methylation over time. (a) Venn diagram of DMGs identified in plants under combined stress (ST4_1_3), and during physiological recovery (ST6_1_3) in season 1, and in primed plants (ST4□00vsST4□33) under a triggering stress. Inserted table represents the direction of changes in methylation (↑ hypermethylation, ↓ hypomethylation) and the functional annotation of 20 common differentially methylated genes across all three time points. (b) Heatmap of level of methylation changes for DMGs that were differentially methylated in ST4_1_3, ST6_1_3, and ST4_2_00 vs. ST4_2_33. Red: hypermethylation. Blue: hypomethylation. Yellow: not differentially methylated.

### The potential relationship between DNA methylation and gene expression

For a better understanding of the potential functional role of DNA methylation on gene expression, we focused on the DMRs overlapping with gene promoters and gene bodies (DMGs). The relationship between changes in gene methylation and transcriptional changes was examined (Figure 5A, B, top). For ST4_2_33 plants, 15 DEGs with DMGs located in the promoters and 8 DEGs with DMGs located in the gene body were identified. Among the 15 located in the promoter, 5 were found to be hypermethylated and down-regulated, 5 were hypomethylated and down-regulated, and 5 were hypomethylated and up-regulated. Of those located in the gene body, 4 were found to be hypermethylated and down-regulated, 1 was hypermethylated and up-regulated, 2 were hypomethylated and down-regulated and 1 was hypomethylated and up-regulated (Figure 5A, bottom). In ST4_2_03 plants, 11 DEGs with DMGs located in the promoter and 12 DEGs with DMGs located in the gene body were identified. Among the 11 located in the promoter, 4 were hypermethylated and down-regulated, 2 were hypermethylated and up-regulated, 3 were hypomethylated and down-regulated, while 2 were hypomethylated and up-regulated. For the 11 located in the gene body, 6 were hypermethylated and down-regulated, 2 were hypermethylated and up-regulated, 3 were hypomethylated and down-regulated, while 1 was hypomethylated and up-regulated (Figure 5B, bottom). No overlapping DEGs and DMGs were identified in ST4_2_30 plants.

**Figure 5.**
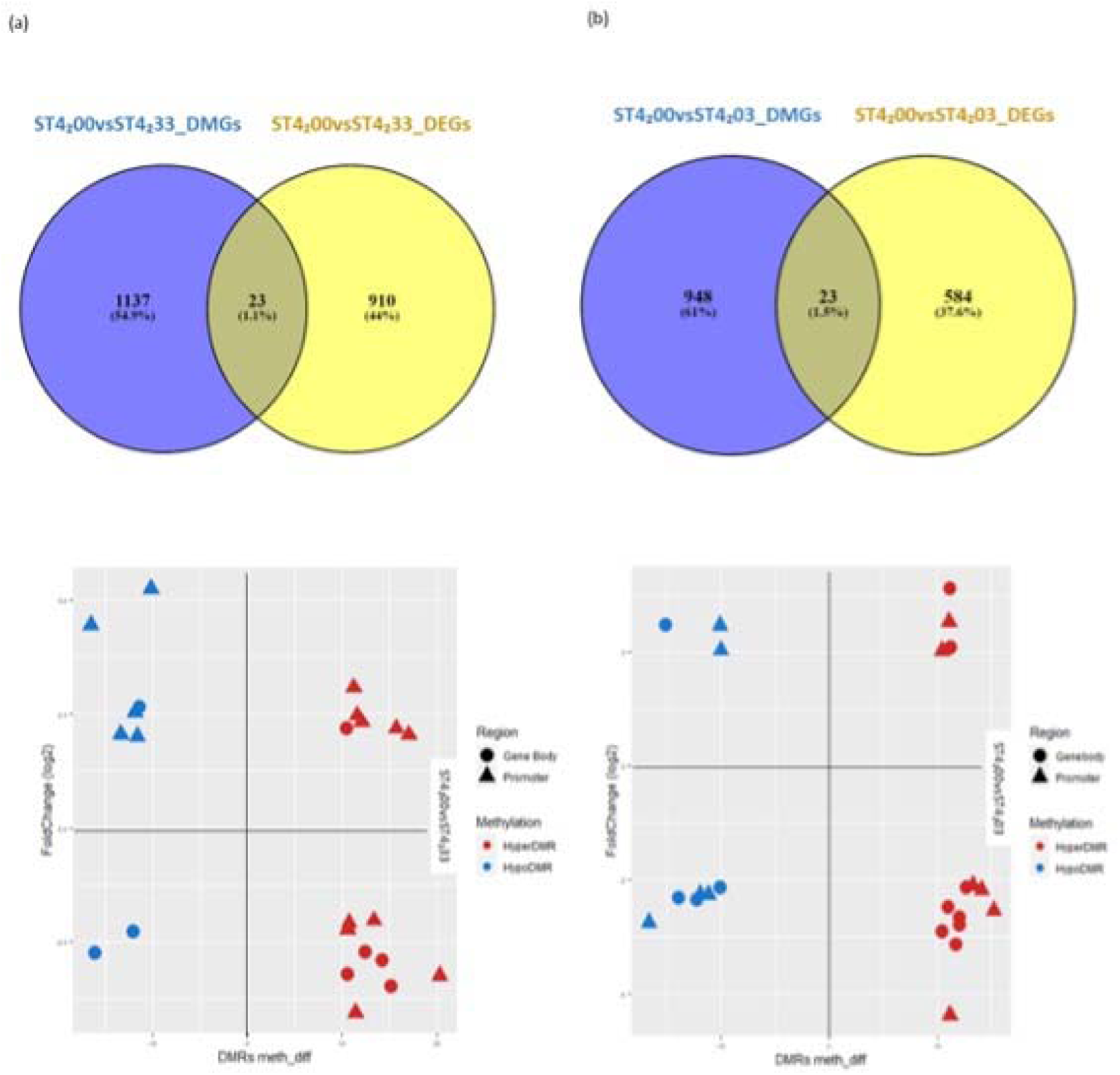
Graph representation of overlapping DEGs and DMGs based on group comparison. (a) Top: venn diagram of overlapping genes between DEGs and DMGs in ST4_2_00 vs ST4_2_33. Bottom: scatterplot of DEGs related with DMGs that located in promoter and gene body, showing the relationship between transcript levels (fold change: log2) and DNA methylation (meth_diff) in ST4_2_00 vs ST4_2_33. (b) Top: venn diagram of overlapping genes between DEGs and DMGs in ST4_2_00 vs ST4_2_03. Bottom: scatterplot of DEGs related with DMGs that located in promoter and gene body) showing the relationship between transcript levels (fold change: log2) and DNA methylation (meth_diff) in ST4_2_00 vs ST4_2_03.

In general, the presence of hypo- or hyperDMGs did not correlate with the transcriptional pattern (up- or down-regulation) of genes. Gene functional annotation analysis revealed the involvement of sHSPs in both ST4_2_03 and ST4_2_33. Regardless of their methylation pattern, these sHSPs were all up-regulated (Table S6-8). Other genes associated with basic Helix-Loop-Helix (bHLH), NAC, B3 transcription factor family genes, as well as SUI1-domain, J-domain, tumor overexpressed genes (TOG)-domain, PMR5N domain, and WD-repeats-region domain-containing protein were identified (Table S7-8).

## Discussion

### Modified response in gene expression after priming

Plants are able to establish a molecular memory of environmental stress (priming stress) that results in an enhanced response to subsequent stresses (triggering stress) (Crisp et al., 2016). Multiple studies suggest that the molecular basis of such memory is epigenetic in nature (Lämke et al., 2016; Liu et al., 2018). Most of this research has been conducted in annual plants, and the few examples of epigenetic memory of stress research in perennials studied epigenetic memory over short periods (days) between the priming and the triggering stress, and none of these studied the effect of winter dormancy on the maintenance of such memory. Our previous work showed that grapevines exposed to combined heat and drought stress induced differential expression of genes associated with epigenetic modifications during and after stress removal, and that GO terms associated to response to stress (i.e., starch and sucrose metabolism and pentose and glucuronate interconversion (Liu et al., 2020b) were enriched for differentially expressed genes after physiological recovery, (i.e., 16 days past the removal of the environmental insult) (Tan et al., 2023). These results suggested the potential establishment and maintenance of epigenetic memory of stress in grapevine over multiple weeks within one growing season. Here we studied the effect of a priming stress on the vine response (i.e., changes in gene expression and DNA methylation) to a triggering stress of the same nature, occurring after a long period of no stress (approximately one year) and after winter dormancy, with the ultimate goals of identifying and describing grapevine genes associated with memory of stress and determining the potential contribution of DNA methylation towards its establishment and maintenance.

First, we compared the response observed in naïve plants under combined stress (ST4_2_03) to that observed in plants under the same conditions in season 1. Gene expression results suggested a degree of consistency in naïve plants’ response to stress for the first time, irrespective of the year of exposure, in terms of the number of differentially expressed genes and their function. Similarly, KEGG pathway analysis revealed the involvement of ‘plant hormone signal transduction’, ‘protein processing in endoplasmic reticulum’, ‘pentose and glucuronate interconversions’, and ‘phenylpropanoid biosynthesis pathway’ genes, consistent with the results previously observed and confirming the importance of those pathways and pathway-associated genes in stress response in naïve plants (Tan et al., 2023).

Primed plants under combined stress (ST4_2_33) showed differences in expression compared to the results observed in naïve plants under combined stress (ST4_2_03). One common feature of the primed state is the reprogramming of the primed plant transcriptome. Such reprogramming results in differences in gene expression between naïve and primed plants in different temporal contexts. These include primed plants presenting 1) different transcriptional patterns than naïve plants even in the absence of a triggering stress (Lämke and Bäurle, 2017); 2) different transcriptional patterns in response to a triggering stress (Ding et al., 2012); and 3) significant differences in the scale of expression change in response to the triggering than the priming stress (Lämke and Bäurle, 2017). Our results from season 1 showed that primed plants presented different transcriptomes than naïve plants 16 days after the removal of the priming stress (i.e., at physiological recovery). These primed plants still showed different transcriptome profiles than naïve ones more than 11 months after the priming stress and after winter dormancy with the identification of a small number of DEGs (39) between primed plants (ST4_2_30) and naïve plants (ST4_2_00) in the absence of second stress. unctional annotation of one of the up-regulated DEGs revealed that it was a histone-lysine N-methyltransferase ATXR6-associated gene. ATXR6 has been reported to deposit histone 3 lysine 27 mono-methylation (H3K27me1) (Jacob et al., 2009) to promote heterochromatin formation, which represses transposable elements (TEs), and controls genome stability in Arabidopsis (Ma et al., 2018). Interestingly, ATXR1, a gene from the same protein family, is necessary but not sufficient for transcriptional memory response (Ding et al., 2012). The involvement of transcription factor regulation in stress response has been well-studied, as they are required to reprogram stress-related genes (Ohama et al., 2017). Specific transcription factor families such as AP2/ERF, NAC, WKRY, and ABC identified among the 39 DEGs in ST4_2_30 have been shown to play important roles in response to abiotic stress such as heat and drought in plants (Chen et al., 2012; Hu et al., 2010; Licausi et al., 2013). More importantly, AP2/ERF and NAC families are involved in stress memory (Ding et al., 2014), although the transcriptional memory pattern of a transcription factor does not necessarily determine the memory pattern of its target gene (Ding et al., 2013; Jacques et al., 2021). Our results suggest that these transcription factors and ATXR6 could be contributing to the maintenance of the long-term somatic stress memory in grapevine. Nevertheless, the establishment and maintenance of epigenetic memory of stress would not be of any biological significance if it did not alter the primed plant response to a recurrent stress. We observed that the exposure to a triggering stress led not only to a higher number of differentially expressed genes but also a larger change in expression of those genes commonly expressed between naïve (ST4_2_03) and primed (ST4_2_33) plants.

This modified gene expression, even after a long period (∼1 year) with no exposure to stress, suggested that stress priming-induced somatic memory is long-term and relatively stable in grapevine, contrary to the somatic stress memory found in annual plants which appears to be transient (Feng et al., 2016; Singh et al., 2014). GO analysis showed the heavy involvement of histone methylation and histone-lysine methylation in both naïve and primed plants response to stress (Supplemental Table S2), suggesting the potential role of histone modification in establishing, maintaining, and retrieving this long-term stress-induced somatic memory, as in previous research correlating histone methylation with somatic stress memory (Lämke et al., 2016).

### Identification of putative stress memory genes

The expression patterns of certain genes in this study closely resembled the expression patterns of stress memory genes, which are stress-inducible genes that have been linked to stress memory establishment (Charng et al., 2007; Charng et al., 2006; Ding et al., 2012; Lämke et al., 2016; Liu et al., 2018). Previous studies have used different systems to classify memory genes based on their transcriptional profile. The first system, described in detail in Bäurle (2018), includes three types: type I, type II and non-memory genes. The expression patterns observed in C7_ST4_, C8_ST4_, and C4_ST5_ could potentially be type I memory genes, as characterized by the gene expression that persists through the recovery phase. Whereas C2_ST4_ and C1_ST5_ could potentially be type II memory genes, where the response is modified, and usually stronger and faster during second exposure (Bäurle, 2018). However, several clusters that do not resemble type I, type II, or non-memory gene expression patterns were also identified (C1_ST4_, C3_ST4_, & C4_ST4_; C2_ST5_ & C3_ST5_). They displayed an opposite expression pattern upon the encounter of the first and second stress. In the second classification system, presented in Ding et al. (2014), transcriptional changes are indicated by (+/+), (-/-), (+/-), (-/+), (-/=), and (+/=), where the first symbol indicates the transcriptional changes compared to control, and second symbol indicates the transcriptional changes compared to first stress response. Most of the clusters we identified in grapevine could be put into one of these categories. However, some of the clusters still could not be classified clearly. Genes in both clusters C8_ST4_ and C2_ST5_ showed a change in expression upon first stress exposure with an incomplete return to the original expression state at physiological recovery, and a lack of response upon encounter of a triggering stress. To account for this, we propose a modified system to classify the memory genes identified in our experiments with perennial plants. In general, it follows the type I and type II classification, but also separates the different expression patterns based on the expression changes occurred upon second stress exposure (-/+/=/0). This modified classification system provides a simple and intuitive visual representation of how gene expression of potential memory genes changes upon the encounter of recurring stress signal and during the period of no stress/recovery.

GO and functional annotation analysis suggested that type I memory genes in grapevine are mainly involved in transcription regulator activity, catalytic activity, and binding, and mainly belong to chaperones. Moreover, the majority of type I memory genes are associated with sHSPs. The memory genes that displayed an opposite regulation profile between first and second stress (Type I^-^), belonged to a wide range of groups such as protein modifying enzyme and molecular function regulators. Interestingly, the well-characterized type II memory gene, heat stress transcription factor A-2 (HSAF2) associated gene (Charng et al., 2007) ortholog (VIT_04s0008g01110) in grapevine was found as part of the C7_ST4_, which contains Type I^=^ memory genes, suggesting that the expression pattern of memory genes may differ in perennial and annual plants.

Type II memory genes identified here, mainly belong to transporter, gene-specific transcriptional regulator, transmembrane signal receptor and chromatin/chromatin binding, or regulatory proteins. A number of chromatin -regulating enzymes and transcription factor associated genes were identified as type II memory genes. In particular, GO analysis of C1_ST5_ (containing type II^+^ memory genes) showed enrichment in methylation, including histone H3-K9 methylation and DNA methylation, suggesting a potential role of epigenetic chromatin based mechanisms in the regulation and maintenance of stress memory genes as in (Lämke and Bäurle, 2017).

### Alteration of DNA methylation patterns under combined stress

A significant increase in global DNA methylation was observed only in primed plants under combined stress (ST4_2_33) (Figure 3A). Previous studies have reported a loss of global DNA methylation under heat stress (Li et al., 2016), while an increase in global DNA methylation is associated with drought stress in more drought-tolerant maize (Wang et al., 2021). The PCA analysis showed that the global DNA methylation pattern of our plants under controlled conditions (ST4_2_00 and ST4_2_30), regardless of priming status, appeared to be variable. while the global DNA methylation pattern of plants under stressed conditions (ST4_2_03 and ST4_2_33) was more conserved (Figure S4). This suggests that cytosine methylation may have arisen stochastically under control conditions, whereas stress-induced DNA methylation was non-random (Feiner et al., 2022). Interestingly, when performing the DMC analyses, more DMCs were identified in the CHH context for primed plants under combined stress, while more DMCs in CG and CHG contexts were identified for both naïve plants under combined stress (ST4_2_03) and primed plants under control conditions (ST4_2_30) (Figure 3B). Methylated CHG and CHH are typically found in silenced regions of the genome such as transposons and repeats, whereas methylated CG may be associated with gene expression regulation (Cokus et al., 2008). The high number of DMCs found in CHH context for primed plants under combined stress might be associated with the modified response to stress triggered by stress memory.

For DNA methylation changes over time, only a small number of DMGs were commonly differentially methylated in our two-year experiment (Figure 4). The limited overlapping DMGs and lack of consistent patterns suggests that DNA methylation might have been reset after stress was over.

### Stress-induced transcriptional regulation is partially independent of DNA methylation

Despite the general belief that DNA methylation in the promoter region of genes inhibits gene expression by influencing the binding of transcriptional activators or repressors (Zhang, Lang & Zhu, 2018), and gene body methylation (GbM) is positively correlated with expression (Yang et al., 2014), we did not observe this correlation between DNA methylation changes (hyper- or hypomethylation) in different genic regions (gene body or promoter) and transcriptional patterns (up- or down-regulation) of the overlapping genes. Moreover, only around 2% of the DMGs were differentially expressed (23/1160, and 23/971 for ST4_2_00vsST4_2_33 and ST4_2_00vsST4_2_03, respectively) (Figure 5). A similar portion and no correlation between methylation changes and transcriptional patterns has been observed in previous studies (López et al., 2022; Rambani et al., 2020). Taken together, our results suggest that stress-induced transcriptional regulation might be, at least partially, independent of DNA methylation. This result is similar to what has been observed in tomatoes for the flower-to-fruit transition, where the variation in the expression of the majority of genes was associated with a change in histone mark distribution, only a minor fraction of differentially expressed genes were associated with DNA methylation (Hu et al., 2021).

Although the involvement of transcription factors in transcriptional memory has been characterized (Ding et al., 2014; Jacques et al., 2021), whether there is an epigenetic basis for such involvement has seldom been studied. In our study, we found that a small number of differentially expressed sHSPs and transcription factors were also differentially methylated. DNA methylation may play a role in regulating the expression of those genes, which then contributes to somatic stress memory, however, due to the overwhelmingly small proportion (2%) of DEGs that were also DMGs, we cannot determine here whether this association was random or not. Therefore, it might be safe to say that the establishment, maintenance, and retrieval of stress-induced long-term somatic memory in grapevine through priming appeared to require more than DNA methylation alone. Histone modification might be the key player in these processes. Our data correspond with others that show that the reprogramming of genes was correlated with their histone marking status but not with changes in cytosine methylation, indicating that histone posttranscriptional modifications rather than DNA methylation are associated with the remodeling of the epigenetic landscape (Hu et al., 2021). Although we did not address histone modification explicitly, the presence of many stressed-induced DEGs (primed or naïve) that are histone/chromatin modification-associated was an indication of the importance of histone modification, consistent with previous research where specific histone modification marks have been shown to not only play a role in drought memory establishment and retrieval but also in heat and salinity stress memory (Ding et al., 2012; Lämke et al., 2016; Sani et al., 2013).

## Conclusions

Plant priming, and subsequent stress memory establishment, maintenance, and retrieval are seldom studied in woody perennial species. Our two-growth season study demonstrated the establishment of molecular stress memory in grapevine and that the established somatic memory could be maintained through winter dormancy. This memory primed the grapevine for a modified transcriptional response upon encountering a second stress, reflected in more DEGs and the magnitude of the change in expression. We identified potential key factors, such as sHSPs and transcription factor families AP2/ERF and NAC in the maintenance of this somatic memory. We also identified and characterized potential stress memory genes in grapevine based on their transcription patterns. In addition to the modified transcriptional response, we observed an increase in global DNA methylation for primed plants compared with naïve plants, and the global DNA methylation profile appeared to be more variable for plants under controlled conditions compared to plants under combined stress, regardless of their priming status. The lack of consistent methylation pattern and small number of overlapping differentially methylated genes before and after stress and after winter dormancy suggests that DNA methylation induced by stress varies and is largely reset between each stress encounter for drought and heat response in grapevine. We also observed that changes in DNA methylation and gene expression do not necessarily coincide with second stress exposure. This suggests that stress memory establishment, maintenance, and retrieval might be more complex and involve multiple epigenetic mechanisms such as histone modification. It remains to be tested if such epigenetic changes can be inherited during clonal propagation, which is common in grapevine, and if such changes could contribute to adaptation to changing environments.

## Materials and methods

### Plant materials and experimental design

To test the establishment, maintenance, and priming effect of long-term memory of stress in *V. vinifera,* a two-growing seasons experiment was carried out during 2016, 2017, and 2018 (Figure 6). Plant material and growth conditions during the first growing season are described in detail in Tan et al. (2023b). In short, 64 propagated dormant cuttings obtained from six donor vines (*V. vinifera* L. Cabernet Sauvignon) were randomly allocated into two different groups (i.e., control and combined drought and heat stress (T0 and T3 respectively hereafter)), and randomly divided into five replicate plots. Plants were then exposed to combined drought and heat stress as described in Tan et al. (2023b). Briefly, the experimental design incorporated drought and high temperature stresses in a factorial design as in Edwards et al (2011). First, irrigation was removed from the selected plants (T3) until they were under moderate to severe drought stress (i.e., stomatal conductance to water vapor (gs), measured using a Delta-T AP4 Porometer (MEA, Magill, SA, Australia), between 75 and 100 mmol/m2/s. Once plants reached this stage, pots were individually weighed and subsequently daily hand-watered to this weight for the duration of the treatment. After 10 days of drought stress, heat stress was applied to selected plants (T3) for 48 hours, by allowing insolation to heat the chamber to 45°C during the day and at a minimum of 30°C during the night. After stress treatment, all plants were maintained under control greenhouse conditions and left to enter winter dormancy at the end of the 2016/17 season. Post-leaf fall, the vines were pruned to a single cane with four buds from the origin on the main stem. Prior to the spring of the second growing season, plants from each of the treatments (0, naïve plants hereafter; and 3, primed plants hereafter) were randomly assigned to two treatments (control or combined stress) and one of four blocks (each containing 4 groups of 4 plants randomly distributed within the block). This resulted in four groups depending on the first and second season groups: 0,0 refers to naïve plants grown under control conditions in season 2; 0,3 refers to naïve plants grown under combined stress in season 2; 3,0 refers to primed plants grown under control conditions on season 2; and 3,3 refers to primed plants grown under combined stress in season 2 (Figure 6A). Each block was placed on a separate bench in a glasshouse (CSIRO, Waite Campus, Adelaide, South Australia, Australia) maintained at an air temperature of 27°C Day/20°C Night, until stress treatments were applied. Humidity and light were uncontrolled. Air temperature and humidity were continuously recorded using a TinyTag Plus 2 logger in a small Stephenson shield (Hastings Data Loggers, Port Macquarie, NSW, Australia).

**Figure 6.**
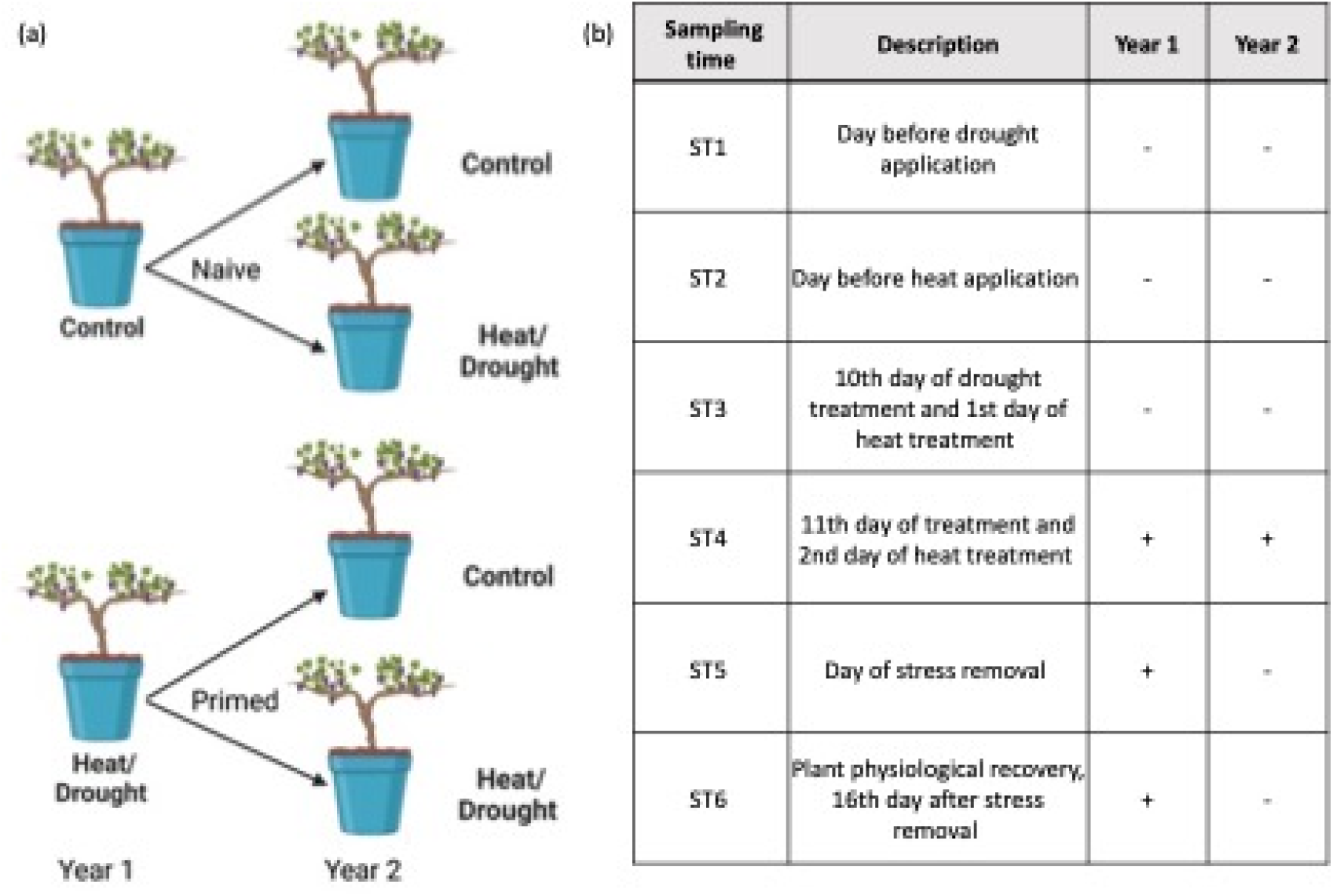
Experimental design. (a) Scheme of stress treatment time course and plant assignment over two growing seasons. (b) Sampling and data collection times. Leaf samples collected at the time points indicated with a + were used for nucleic acid extractions to analyze gene expression and DNA methylation differences between plants grown under control and stress conditions.

Water stress was imposed by removing drippers and monitoring stress using measurements of stomatal conductance (g_s_), with small amounts of additional water provided as required to maintain g_s_ in the 50-75 mmol/m^-2^/s^-1^ range. The water stress treatment started on 23/1/2018, with heat stress generated for two days using natural insolation, as in Tan et al. (2023b), on 4-5/2/2018. On the second day of combined stress of year 2, g_s_ was measured on every vine (64 in total) at approximately 4 PM, then a single set of measurements (stem water potential (Ψs), leaf temperature, g_s_ and leaf sampling (snap-frozen leaves for DNA and RNA analysis) was undertaken (Figure 6B), as described in Tan et al. (2023b), with the exception that it was not possible to take g_s_ measurements on every leaf (2-3 reps per 2017/2018 treatment).

The second and third leaves down from the apical meristem were sampled at four time points during the two seasons (Figure 6B). Four replicates were collected from each sampling time/treatment combination in season 1, and eight replicates were collected from each sampling time/treatment combination in season 2. Samples were coded according to their sampling time (ST4, ST5, or ST6), season (1 or 2) and treatment (control (0) or heat and drought (3)). Season 1 samples are described using a five-character code, i.e., leaf samples collected at sampling time 4 of season 1 from plants under control conditions were coded ST4_1_0, while samples from season 2 are described using six characters, i.e., a sample collected at sampling time 4 of season 2 from naïve plants under control conditions were coded ST4_2_00.

### Nucleic acid extraction

Sample leaves from each plant were frozen immediately after collection using liquid nitrogen and stored at -80°C. Frozen leaves were ground to a fine powder under liquid nitrogen using mortar and pestle. Samples were split into two subsamples and stored at -80°C until further use.

Total RNA was extracted from 100 mg of frozen and ground samples using the Spectrum™ Plant Total RNA Kit (Sigma, St. Louis, Missouri, USA) according to the manufacturer’s Protocol A. Spectrophotometric analysis (NanoDrop™ 1000, Thermo Fisher Scientific, Wilmington, DE, USA) and Experion™ RNA StdSens Chips (BIO-RAD, USA) were used to determine RNA integrity. Only samples with a RNA quality indicator (RQI) above 7 and presenting 260/280 and 260/230 absorbance ratios between 1.8-2.2 were used for library preparation. 4 μg of total RNA per sample was used for ribosomal RNA depletion using Dynabeads mRNA Purification Kit (Thermo Fisher Scientific, USA) following the manufacturer’s instructions.

Total DNA was extracted from 100 mg of frozen ground samples, using the DNeasy Plant kit (Qiagen). The concentration and integrity of the DNA were measured by Fragment Analyzer (Agilent Technologies).

### RNA Sequencing (RNASeq)

5 μl of ribosomal depleted RNA from each sample were used to prepare 64 individually barcoded RNA-seq libraries using the NEBNext® Ultra™ RNA Library Prep Kit for Illumina (New England Biolabs, USA) following the manufacturer’s instructions. The Illumina NextSeq 500 HighOutPut platform was used to produce 75bp single-end runs at the Australian Genome Research Facility (AGRF) in Adelaide, Australia.

### Whole Methylome Sequencing (WMS)

Whole Methylome Sequencing (WMS) was performed on genomic libraries prepared using the NEBNext Enzymatic Methyl-seq Kit (New England BioLabs) following the manufacturer instructions. Each individual sample of genomic DNA was spiked with internal controls to determine the enzymatic conversion efficiencies and the abundance of false positives and negatives (i.e., 0% methylated Lambda DNA, and 100% CpG methylated pUC19 DNA). Spiked DNA samples were then fragmented to 200 – 300 bp using the Covaris S220 ultrasonicator. The resulting individually barcoded libraries were sequenced using Nova Seq 6000, and PE150 with a paired-end sequencing approach.

### Bioinformatics Analyses

#### RNA-sequencing data analysis

Raw sequencing data were processed on the LipsComb Compute Cluster (LCC) platform at the University of Kentucky, United States. AdapterRemoval (Lindgreen, 2012) was used for removing adaptors of the raw reads. Sequence quality control was performed with FastQC (http://www.bioinformatics.babraham.ac.uk/projects/fastqc/) (2015). The reads were mapped to a 12X grapevine reference genome (NCBI assembly ID: GCF_000003745.3) with the alignment tool (HISAT2) (Kim et al., 2015). The GTF reference of the *Vitis vinifera* genome was downloaded from the *Ensembl Plants* website (http://plants.ensembl.org/Vitis_vinifera/Info/Index). Samtools (Li et al., 2011) was used to generate Binary Alignment Map (BAM) files after mapping the reads to the genome.

#### Identification of putative memory genes using co-expressed gene cluster analysis

Transcripts Per Million (TPM) of each plant sample were calculated from the BAM files using the TPMcalculator (Alvarez et al., 2019). Normalized data (calculated TPMs, log2 transformed) was used for the identification of gene expression clusters based on gene expression patterns during the following time point/treatment combinations: control plants sampled in season 1 ST4 (ST4_1_0), stressed plants sampled in season 1 ST4, ST5, and ST6 (ST4_1_3, ST5_1_3, and ST6_1_3), and primed plants under combined stress sampled at season 2 ST4 (ST4_2_33) using *clust* v1.8.4 (Abu-Jamous & Kelly, 2018). Resultant clusters were then classified according to three conditions: A) If the gene expression level in stressed plants at physiological recovery was significantly different than that presented by control plants (ST4_1_0 ± ST6_1_3; T-Test p-val < 0.05) or not, b) if the change in expression in response to the triggering stress was significantly different than in response to the priming stress (ST4_1_3 ± ST4_2_33; T-Test p-val < 0.05), and c) if the triggering stress induced a significantly different change in expression compared to the expression level at physiological recovery from the priming stress (ST6_1_3 ± ST4_2_33; T-Test p-val < 0.05).

#### Differentially expressed genes (DEGs) analysis

Gene expression was estimated using the *edgeR* package (Robinson *et al*., 2010) on Rstudio. The raw mapped data of each sample was standardized by edgeR’s trimmed mean of M values (TMM). This method estimates scale factors between samples to determine DEGs. Between control and treatment, a log2fold change(log2FC) of 2 and a false discovery rate adjusted P-value<0.05 using Benjamini and Hochberg’s algorithm was adopted to indicate significance. This process was repeated for each group of comparisons.

#### Gene ontology (GO), DEGs visualization, and functional annotation

All differentially expressed genes of interest were subjected to ontology analysis through using of agriGO v2.0 (Tian et al., 2017). DEGs of each treatment were used to attain the significant GO terms with agriGO v2.0 with the following criteria: Fisher’s statistical test method, Yekutieli (FDR under dependency) multi-test adjustment method, significance level <0.05, and selecting either complete GO or slim GO as the gene ontology type. The visualization of the expression level of selected DEGs was done through the built-in plot function of R. Functional annotation of DEGs was obtained from PantherDB (Mi et al., 2021). Plots were made with the R-package ggplot2.

#### Identification of differentially methylated cytosines and regions (DMCs and DMRs)

Adaptor sequences, low-quality reads, and contaminants were removed from WMS reads using Adapter Removal V2 software. The enzymatic conversion efficiency of unmethylated and methylated cytosines was calculated using pipelines (https://github.com/nebiolabs/EM-seq/blob/master/em-seq.nf) and the methylation control sequences (https://github.com/nebiolabs/EM-seq/blob/master/methylation_controls.fa) provided by the NEBNext Enzymatic Methyl-seq Kit manufacturer.

Genome indexing was performed with Bismark using ‘--bismark_genome_preparation’ option (Krueger and Andrews, 2011) using the C-to-T and G-to-A versions of the reference grapevine genome (PN40024 v.4) created with Bowtie2 (Langmead and Salzberg, 2012). Sequencing coverage and depth were estimated using Samtools coverage and depth toolkits (Li et al., 2009). Methylation calling was performed with Bismark extractor (https://www.bioinformatics.babraham.ac.uk/projects/bismark/Bismark_User_Guide.pdf) by calling ‘--comprehensive’ and ‘--cytosine_report’ option after the conversion to bedGraph. Both Differentially methylated cytosines and regions (DMCs and DMRs respectively) were determined using the ‘Methylkit’ package (Akalin et al., 2012) with default parameters (minimum coverage threshold of 10 and 5 for DMCs and DMRs, respectively; q-value ≤ 0.05; minimum differential methylation level of 10%); sliding window for DMRs was 1000 bp). Genes were deemed differentially methylated when a DMR overlapped with their promoter (defined here as 1000 bp upstream of the transcription starting site (TSS)), or with the body of the given gene.

## Author Contributions

CML, PT and EJE conceived and designed the study. YH and KT perfomed the greenhouse and laboratory experiments. JWT and HS performed the analysis and analyzed the results. All listed authors contributed to the manuscript substantially and have agreed to thre final submitted version.

## Funding

This study was supported by the Australian Grape and Wine Authority grant ID: UA1503, the National Institute of Food and Agriculture, AFRI Competitive Grant Program Accession number 1018617, and the National Institute of Food and Agriculture, United States Department of Agriculture, Hatch Program accession number 1020852.

## Conflict of Interest

Author Penny Tricker is employed by The New Zealand Institute for Plant and Food Research Limited, author Everard J. Edwards is employed by CSIRO Agriculture & Food. The remaining authors declare that the research was conducted in the absence of any commercial or financial relationships that could be construed as a potential conflict of interest.

## Data Availability

The RNA sequencing reads of all plants from first and second growing season were deposited in the sequence read archive (SRA) of the National Center of Biotechnology Information (NCBI) under the accession number PRJNA662522 and , respectively. The DNA sequencing reads of all plants from first and second growing seasons were deposited under the accession number.

