## Supplemental Figure S1 for "Transcriptome Analysis Reveals Long-Term Somatic Memory of Stress in The Woody Perennial Crop, Grapevine *Vitis Vinifera* cv. Cabernet Sauvignon"

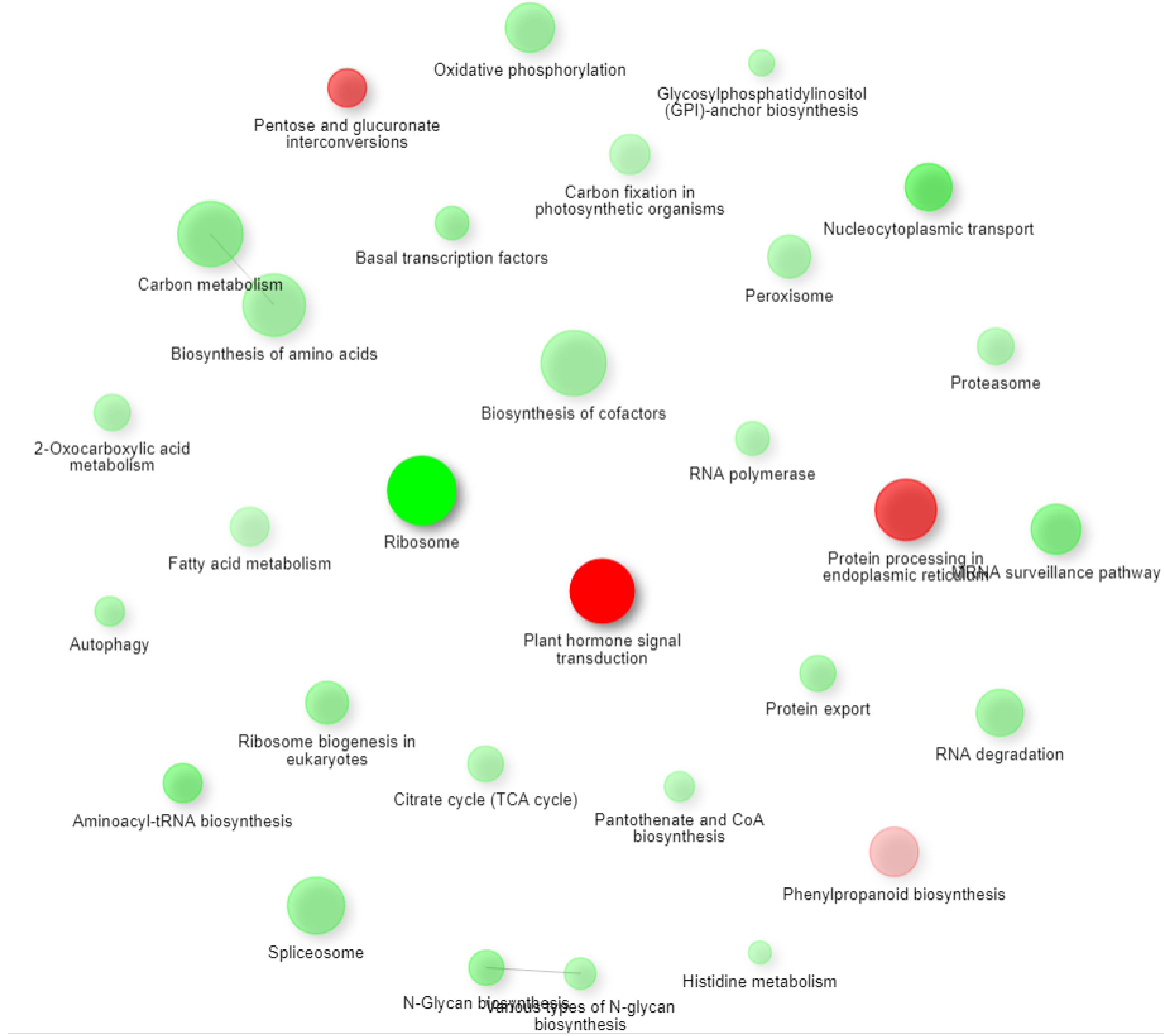


**Supplemental Figure S1. KEGG pathway analysis of DEGs identified in T00vs T03.** Red: up-regulated DEGs. Green: down-regulated DEGs. p-adj $\leq$ 0.05 was used to determined significance.
