## Supplemental Figure S2 for "Transcriptome Analysis Reveals Long-Term Somatic Memory of Stress in The Woody Perennial Crop, Grapevine *Vitis Vinifera* cv. Cabernet Sauvignon"

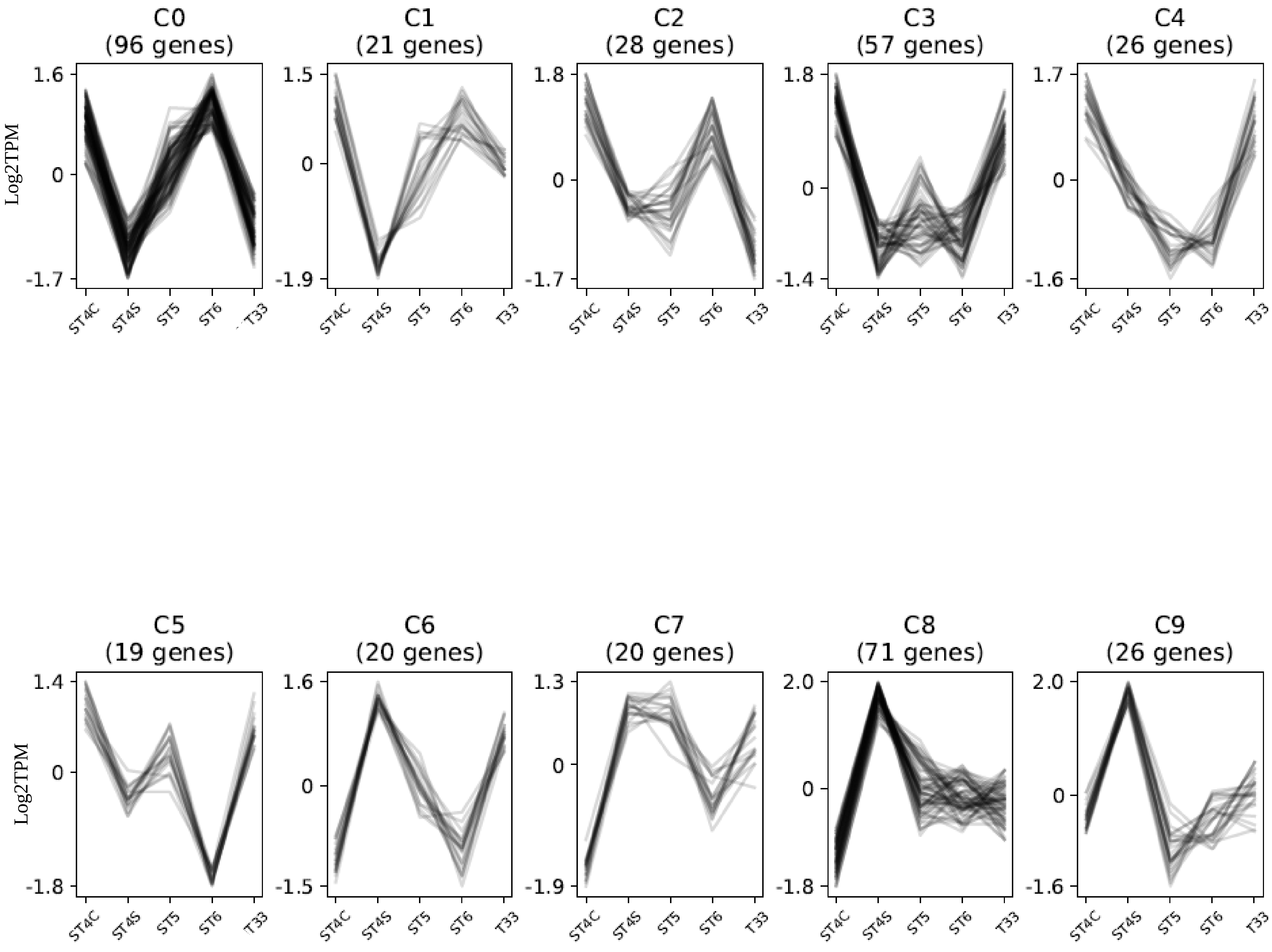


**Supplemental Figure S2. Clusters of DEGs from ST4 sampling time point of first year.** ST4C: Control plants during 12th day of drought and 2nd day of heat stress event (ST4). ST4S: ST4 combined stress plants. ST5: Combined stressed plants at stress removal (ST5). ST6: Combined stress plants during physiological recovery. T33: Primed plants under combined stress.
