## Supplemental Figure S4 for "Transcriptome Analysis Reveals Long-Term Somatic Memory of Stress in The Woody Perennial Crop, Grapevine *Vitis Vinifera* cv. Cabernet Sauvignon"

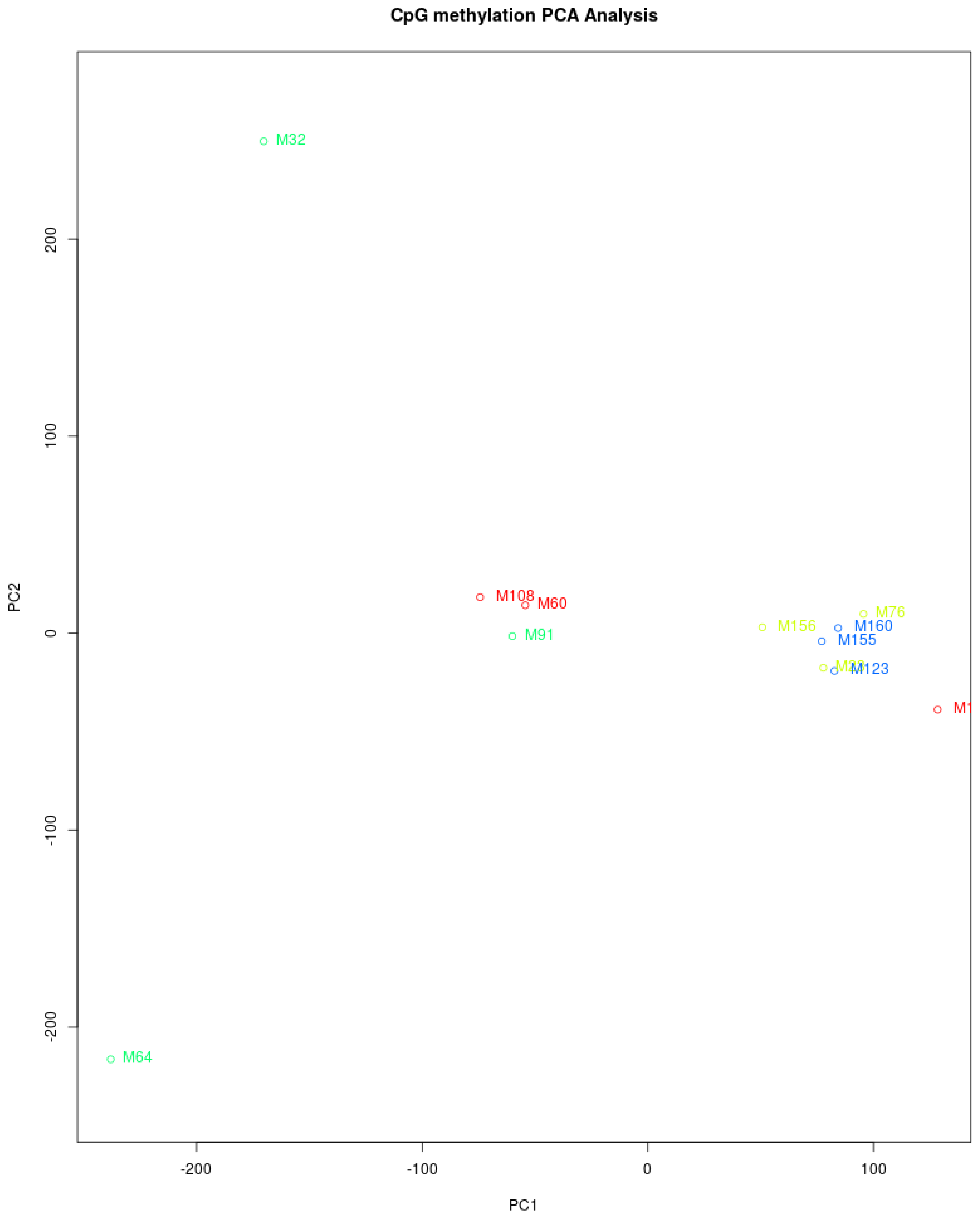


**Supplemental Figure S4. Principle Component Analysis (PCA) plot of the plant methylation patterns separated by their treatments.** Red: T00 plants. Green: T30 plants. Yellow: T03 plants. Blue: T33 plants.
