## Supplemental Table S1 for "Transcriptome Analysis Reveals Long-Term Somatic Memory of Stress in The Woody Perennial Crop, Grapevine *Vitis Vinifera* cv. Cabernet Sauvignon"

| Plant ID | Total Sequence Reads | alignment % |
|---|---|---|
| **P13M** | 28618060 | 80.79% |
| **P55M** | 27672452 | 77.85% |
| **P60M** | 23717195 | 77.78% |
| **P104M** | 22419319 | 79.64% |
| **P107M** | 26397032 | 79.34% |
| **P108M** | 22458021 | 78.80% |
| **P111M** | 25132491 | 79.81% |
| **P112M** | 23937167 | 80.41% |
| **P139M** | 24558296 | 80.94% |
| **P23M** | 30010997 | 87.86% |
| **P24M** | 11856548 | 84.33% |
| **P48M** | 28904203 | 88.08% |
| **P51M** | 26833161 | 88.78% |
| **P71M** | 27094568 | 83.13% |
| **P75M** | 29819640 | 86.23% |
| **P155M** | 22757120 | 87.20% |
| **P76M** | 28253133 | 86.50% |
| **P80M** | 25086346 | 89.38% |
| **P91M** | 31390344 | 85.63% |
| **P135M** | 32331815 | 87.37% |
| **P140M** | 27992565 | 86.86% |
| **P148M** | 27916825 | 87.67% |
| **P17M** | 25701448 | 82.06% |
| **P19M** | 21575689 | 79.01% |
| **P31M** | 25868875 | 73.48% |
| **P56M** | 26791247 | 85.02% |
| **P63M** | 29980087 | 79.59% |
| **P100M** | 25317810 | 84.12% |
| **P103M** | 24102044 | 86.57% |
| **P115M** | 28577258 | 79.72% |
| **P116M** | 27037084 | 87.57% |
| **P144M** | 27328761 | 87.75% |
| **P28M** | 25012880 | 66.27% |
| **P47M** | 23450785 | 77.10% |
| **P52M** | 24118792 | 73.38% |
| **P68M** | 26517071 | 79.41% |
| **P88M** | 29638562 | 73.53% |
| **P92M** | 24919778 | 75.99% |
| **P99M** | 31964273 | 72.54% |

| | | |
|---|---|---|
| **P127M** | 28121889 | 73.71% |
| **P128M** | 22543915 | 82.19% |
| **P152M** | 20968244 | 77% |
| **P27M** | 21444376 | 62.98% |
| **P43M** | 20994249 | 82.78% |
| **P119M** | 21649710 | 78.04% |
| **P123M** | 25308571 | 85.05% |
| **P131M** | 24413873 | 65.24% |
| **P151M** | 21565336 | 80.92% |
| **P159M** | 17075729 | 86.70% |
| **P32M** | 23275600 | 78.39% |
| **P72M** | 20012731 | 84.64% |
| **P79M** | 24911448 | 90.66% |
| **P84M** | 25574231 | 69.79% |
| **P143M** | 24814193 | 80.73% |
| **P156M** | 27304255 | 85.02% |
| **P160M** | 25217806 | 84.73% |
| **P11M** | 23255663 | 90.35% |
| **P35M** | 40004448 | 80.49% |
| **P36M** | 24397534 | 82.07% |
| **P59M** | 27094493 | 79.25% |
| **P64M** | 20445685 | 77.69% |
| **P83M** | 26175474 | 84.30% |
| **P132M** | 21250802 | 77.26% |
| **P136M** | 20792888 | 91.60% |

**Supplemental Table S1.** List of plant samples collected for RNA sequencing. Table includes the total RNA sequencing reads generated from each plant sample and the percentage of alignment rate to the reference genome.
