## Supplemental Table S5 for "Transcriptome Analysis Reveals Long-Term Somatic Memory of Stress in The Woody Perennial Crop, Grapevine *Vitis Vinifera* cv. Cabernet Sauvignon"

| | hit | total | alignment % | avg_depth | avg_coverage | Bisulfite Conversion rate | Non-conversion rate |
|---|---|---|---|---|---|---|---|
| **M108** | 27992521 | 51206002 | 54.66648 | 13.4395 | 80.3002 | 93.2 | 0.2 |
| **M123** | 36659056 | 67110249 | 54.62512 | 16.6686 | 81.3689 | 96.1 | 0.5 |
| **M140** | 26705889 | 50114706 | 53.28953 | 12.9874 | 80.3671 | 97 | 0.4 |
| **M155** | 58459820 | 107242183 | 54.51196 | 25.369 | 83.1472 | 94.8 | 0.2 |
| **M156** | 35415280 | 65933622 | 53.71354 | 16.0133 | 81.3714 | 96.7 | 0.2 |
| **M160** | 31601674 | 57263833 | 55.1861 | 15.0108 | 80.7841 | 94.9 | 0.2 |
| **M23** | 46898885 | 89228145 | 52.56064 | 21.0965 | 82.3533 | 97.5 | 0.2 |
| **M32** | 27806018 | 51146853 | 54.36506 | 13.1863 | 80.1009 | 93.4 | 0.2 |
| **M64** | 28736760 | 52254059 | 54.99431 | 12.8465 | 79.7338 | 95.4 | 0.2 |
| **M76** | 30434109 | 55739355 | 54.60076 | 14.1226 | 80.5556 | 96 | 0.2 |
| **M91** | 36411474 | 67103877 | 54.26136 | 16.9364 | 81.2395 | 94.2 | 0.1 |
| **M60** | 58490908 | 110129306 | 53.11112 | 27.305 | 83.3508 | 93.9 | 0.3 |

**Supplemental Table S5.** Table of total methylation reads, alignment percentage to the reference genome, average sequencing depth, average genome coverage, the average bisulfite conversion rate, and the non-bisulfite conversion rate by sample.
