## Supplemental Table S7 for "Transcriptome Analysis Reveals Long-Term Somatic Memory of Stress in The Woody Perennial Crop, Grapevine *Vitis Vinifera* cv. Cabernet Sauvignon"

| | meth_diff | fold change (log2) | promoter | Functional Annotation |
|---|---|---|---|---|
| **VIT_06s0004g03110** | 20.32915 | -3.23938 | 1 | uncharacterized protein (mediator of DNA damage checkpoint protein 1-like) |
| **VIT_06s0080g01230** | 17.07937 | 2.035223 | 1 | uncharacterized protein (DNAJ-like protein) |
| **VIT_13s0156g00130** | 15.73358 | 2.177897 | 1 | uncharacterized protein |
| **VIT_05s0020g00620** | 15.15005 | -3.47441 | 0 | Kinesin motor domain-containing protein (Kinesin-like protein kin-12C) |
| **VIT_12s0059g01070** | 14.21199 | -2.89267 | 0 | Condensin complex subunit 2 |
| **VIT_18s0001g13540** | 13.34696 | -2.03462 | 1 | uncharacterized protein (WAS/WASL-interacting protein family member 1) |
| **VIT_14s0108g00710** | 12.44134 | -2.70995 | 0 | structural maintenance of chromosomes protein |
| **VIT_17s0000g06000** | 12.08503 | 2.322497 | 1 | BHLH domain containing protein (transcription factor BHLH47 |
| **VIT_04s0008g01500** | 11.60643 | 2.461715 | 1 | SHSP domain containing protein (18.0 KDA class II heat shock protein) |
| **VIT_05s0020g01270** | 11.44308 | -4.05115 | 1 | TOG domain containing protein (Arm repeat superfamily protein) |
| **VIT_04s0008g01570** | 11.24929 | 3.072834 | 1 | SHSP domain containing protein (18.0 KDA class II heat shock protein) |
| **VIT_08s0007g06460** | 10.73195 | -2.08921 | 1 | Laccase (Laccase-11) |
| **VIT_01s0137g00360** | 10.61423 | -2.23017 | 1 | WD_repeats_region domain-containing protein (F21J9.19) |
| **VIT_03s0063g00620** | 10.51584 | -3.19493 | 0 | TF-B3 domain-containing protein (B3 domain containing protein OS06G0194400) |
| **VIT_19s0014g01830** | 10.47465 | 2.1874 | 0 | Glutamate receptor (PBPE domain-containing protein) |
| **VIT_18s0089g01270** | -10.1814 | 5.246983 | 1 | SHSP domain containing protein (22.0 KDA heat shock protein) |
| **VIT_03s0088g00320** | -11.4071 | 2.660499 | 0 | uncharacterized protein (Zinc metallopeptidase EGY3, chloroplastic-related) |
| **VIT_09s0002g00690** | -11.6204 | 2.005235 | 1 | J domain-containing protein |
| **VIT_14s0108g01070** | -11.8766 | 2.530252 | 1 | NAC domain-containing protein (NAC domain-containing protein 59) |
| **VIT_17s0000g07440** | -12.0758 | -2.27574 | 0 | replication protein A subunit (REPA_OB_2 domain-containing protein) |
| **VIT_19s0090g00420** | -13.3881 | 2.046525 | 1 | SUI1 domain-containing protein |
| **VIT_13s0084g00080** | -16.1835 | -2.72743 | 0 | uncharacterized protein (65-KDA microtubule-associated protein 5) |
| **VIT_13s0019g03000** | -16.5499 | 4.453701 | 1 | SHSP domain-containing protein |

**Supplemental Table S7.** The relationship between the presence of DMGs (promoters and gene body) and transcriptional changes (DEGs) in T00vsT33. Functional annotations of those 23 overlapping genes.
