## Supplemental Table S8 for "Transcriptome Analysis Reveals Long-Term Somatic Memory of Stress in The Woody Perennial Crop, Grapevine *Vitis Vinifera* cv. Cabernet Sauvignon"

| | Meth_diff | fold change (log2) | promoter | Functional Annotation |
|---|---|---|---|---|
| **VIT_06s0009g00820** | 15.22915 | -2.53305 | 1 | Kinesin motor domain-containing protein (Kinesin-like protein kin-12E) |
| **VIT_13s0064g01260** | 14.11227 | -2.17843 | 1 | DRT100 like protein (DNA damage-repair/toleration) |
| **VIT_04s0008g01700** | 13.32375 | -2.09481 | 1 | uncharacterized protein (Protein big grain 1-like B) |
| **VIT_06s0004g06300** | 12.70744 | -2.11852 | 1 | Cell division control protein 6 homolog |
| **VIT_18s0122g00980** | 12.04286 | -2.78495 | 1 | Glucan endo-1,3-beta-D-glucosidase |
| **VIT_09s0002g01680** | 12.01157 | -2.64902 | 0 | uncharacterized protein (IQ domain-containing protein IQM5) |
| **VIT_08s0056g00210** | 11.70215 | -3.12421 | 1 | Kinase cdc2 homolog B (Cyclin-dependent kinase 2) |
| **VIT_14s0066g01710** | 11.22054 | 2.08894 | 0 | PMR5N domain-containing protein (protein altered xyloglucan 4-like) |
| **VIT_09s0002g00550** | 11.18936 | -4.36708 | 0 | uncharacterized protein (GDSL esterase/lipase 1-like isoform X1) |
| **VIT_18s0122g01300** | 11.13726 | 3.129071 | 0 | uncharacterized protein |
| **VIT_04s0008g01570** | 11.1165 | 2.529929 | 0 | SHSP domain-containing protein (18.0 KDA class II heat shock protein) |
| **VIT_05s0020g02910** | 10.99561 | -2.47325 | 1 | Protein kinase domain-containing protein (Mitogen-activated protein kinase - kinase kinase 3) |
| **VIT_11s0016g04960** | 10.47104 | 2.02772 | 0 | uncharacterized protein |
| **VIT_15s0021g01380** | 10.45002 | -2.90209 | 1 | Protein kinase domain-containing protein (Cell dibision cycle 7-related protein kinase) |
| **VIT_18s0001g13810** | -10.0096 | 2.034043 | 1 | Auxin-responsive protein (auxin-responsive protein IAA27) |
| **VIT_09s0002g00640** | -10.0832 | 2.466683 | 1 | SHSP domain-containing protein (17.4 KDA class III heat shock protein) |
| **VIT_08s0007g07490** | -10.1145 | -2.12537 | 1 | uncharacterized protein (Protein SINE1) |
| **VIT_05s0062g00480** | -11.1861 | -2.25557 | 0 | Xyloglucan endotransglucosylase/hydrolase (Xyloglucan endotransglucosylase/hydrolase protein 15) |
| **VIT_01s0026g00530** | -11.933 | -2.25126 | 1 | WAT1-related protein |
| **VIT_10s0003g01990** | -12.3016 | -2.33565 | 1 | non-specific serine/threonine protein kinase |
| **VIT_07s0104g00470** | -13.9421 | -2.30765 | 1 | uncharacterized protein (Protein FANTASTIC FOUR 1) |
| **VIT_03s0088g00320** | -15.1734 | 2.486666 | 0 | uncharacterized protein (Zinc metallopeptidase EGY3, chloroplastic-related) |
| **VIT_13s0019g02530** | -16.6903 | -2.75399 | 1 | uncharacterized protein (cucumisin-like isoform X1) |

**Supplemental Table S8.** The relationship between the presence of DMGs (promoters and gene body) and transcriptional changes (DEGs) in T00vsT03. Functional annotations of those 23 overlapping genes.
